# Targeted genome editing in *Nicotiana tabacum* using inducible CRISPR/Cas9 system

**DOI:** 10.1101/2020.03.13.990085

**Authors:** Chong Ren, Yanfei Liu, Xida Wang, Yuchen Guo, Peige Fan, Shaohua Li, Zhenchang Liang

## Abstract

Targeted genome editing has been achieved in multiple plant species using the clustered regularly interspaced short palindromic repeats (CRISPR)-associated protein 9 (CRISPR/Cas9) system, in which the *Cas9* gene is usually driven by constitutive promoters. However, constitutive expression of *Cas9* is not necessary and can be harmful to plant development. In this study, we developed an estrogen-inducible CRISPR/Cas9 system by taking advantage of the chimeric transcription activator XVE and tested the efficacy of this inducible system in *Nicotiana tabacum* by targeting the phytoene desaturase (*NtPDS*) gene, whose mutation resulted in albino phenotypes. Treatment of four independent transgenic lines with exogenous estradiol successfully induced targeted mutagenesis in *NtPDS*. Sanger sequencing assay uncovered the presence of indel mutations (nucleotides insertions or deletions) at the target site as expected, and at least two types of mutations were identified for each line. Transgenic plants with mutated *NtPDS* gene after estradiol treatment exhibited pale green or incomplete albino leaves. Moreover, the expression of *Cas9* in transgenic plants was strongly induced by estradiol treatment. Our results demonstrate the efficacy of XVE-based CRISPR/Cas9 system in *N. tabacum*, and the system reported here promises to be a useful approach for conditional genome editing, which would facilitate the study of genes of interest, especially those developmentally important genes.

## 1. Introduction

Targeted genome editing (TGE) plays a significant role in functional study of genes of interest, trait improvement of crops and development of new cultivars [1,2,3]. Engineered nucleases, such as zinc finger nucleases (ZFNs), transcription activator-like effector nucleases (TALENs) and clustered regularly interspaced short palindromic repeats (CRISPR)-associated protein 9 (CRISPR/Cas9), are generally employed to accomplish TGE in plants [4,5,6,7]. Among them, the CRISPR/Cas9 system is predominantly used for TGE due to its simplicity, high efficiency and versatility [8]. In most cases, the Cas9 encoding gene is driven by constitutive promoters, such as CaMV 35S and ubiquitin promoters [9,10,11,12,13]. However, constitutive expression of *Cas9* might be harmful to cells, or increase the risk of off-target effect [14,15]. More importantly, it is not feasible using this constitutive CRISPR/Cas9 system to generate homologous knockouts of developmentally important genes, especially those genes involved in regeneration, reproduction, or even lethality.

Inducible gene expression systems allowing temporal expression of target genes are employed as powerful tools for research in plant functional genomics [16]. Chemical-inducible systems have been widely used in plants, and inducible systems responding to different chemical inducers, such as estradiol [17,18,19], ethanol [20,21,22,23], glucocorticoid [24,25], ecdysone [26], and tetracycline [27] have been successfully developed. However, some of these systems have limitations during their applications. For instance, application of ethanol would cause toxic effects on treated plants, and unwanted activation of gene expression can be triggered in neighboring plants due to the volatile nature of the inducer when using ethanol-inducible system [28]. The application of glucocorticoid-inducible system, however, was found to cause growth defects in several plant species, including Arabidopsis, tobacco and rice [24,29,30]. The estradiol-inducible system, based on transcriptional activator XVE [18], has been applied to multiple plant species, and no physiological or morphological effect was observed on treated plants [16,18,19,31,32]. Inducible systems are previously used to remove selectable markers or investigate expression patterns of target genes in transgenic plants [19,32,33]. Tang and Liu [34] adopted inducible promoters to drive the expression of *Cas9* or base editors to record stimuli events in both bacteria and mammalian cells. In plants, Tang et al. [35] employed the estrogen-inducible (XVE) promoter to drive the expression of CRISPR/Cas9 reagents in rice. Very recently, inducible genome editing was reported in Arabidopsis based on the estrogen-inducible XVE system [36]. However, targeted genome editing based on inducible systems in other plant species is still less studied.

Here we developed inducible CRISPR/Cas9 system by taking advantage of the estrogen-inducible XVE system, and the phytoene desaturase (*NtPDS*) gene, whose mutation generally results in visible albino phenotypes [11,37], was chosen as a proof-of-concept target for conditional genome editing in the model plant *Nicotiana tabacum*. Prior to the inducible genome editing, we first tested the efficacy of the XVE system by investigating gene expression of *GUS* (β-glucuronidase) in transgenic tobacco plants upon estradiol treatment. GUS staining revealed that XVE-controlled expression of *GUS* reporter gene was induced by exogenous estradiol treatment. Similar result was obtained in estradiol-treated transgenic plants using the *Cas9* gene instead of *GUS* reporter gene. Under normal conditions, the transgenic plants showed none of any *PDS*-defective phenotypes, and no mutation was detected at the target site in *NtPDS* gene. By contrast, after exposure to estradiol, the transgenic plants exhibited etiolation or albino phenotypes. Sanger sequencing assay showed that the *NtPDS* gene was successfully edited after estradiol treatment. The obtained results demonstrate the efficacy of XVE-controlled CRISPR/Cas9 system in *N. tabacum*, and our method described here thereby provide a useful approach for conditional genome editing in plants.

## 2. Results

### 2.1. XVE stringently mediates the expression of GUS gene upon estradiol treatment

The *GUS* gene was amplified and introduced into the XVE vector pER8 [18] to generate the estrogen-inducible expression construct pER-GUS (Figure 1a). The GUS reporter is a reliable and extremely senesitive system that allows histochemical assessment of gene activity in plants [16,38]. After transformation of *N. tabacum*, hygromycin-resistant plants that can develop roots on hygromycin-containing medium (Figure S1) were selected as candidates for PCR identification. Those plants identified with exogenous *GUS* gene were selected as transgenic plants (Figure 1b and Figure S2). Among the 98 tested plants, 21 plants were identified with exogenous T-DNA insertions, with a transformation rate of 21.43% (Figure 1c). These obtained T0 plants were subcultured on Murashige and Skoog (MS) medium and were used for estradiol treatment. Sixteen out of the 21 transgenic plants were randomly selected and divided into two groups and were treated with or without estradiol. Histochemical staining revealed strong GUS activity in pER-GUS-transformed plants after treatment with 20 µM estradiol (Figure 2a,b). Interestingly, GUS staining was only detected in roots and lower leaves, while no GUS staining was observed in upper leaves (Figure 2a and Figure S3). As shown in Figure 2a, strong GUS expression was detected in roots and leaf 1-2 of the estradiol-treated lines. By contrast, no GUS staining was observed in leaves (leaf 5 and 6 of line 7, leaf 7 and 8 of line 8, and leaf 8 and 9 of line 9) that are distal to the roots. From the eight independent transgenic lines without treatment with estradiol, two lines showed weak GUS staining in roots, and the others, however, exhibited no GUS staining in either leaves or roots (Figure 2a,b).

**Figure 1.**
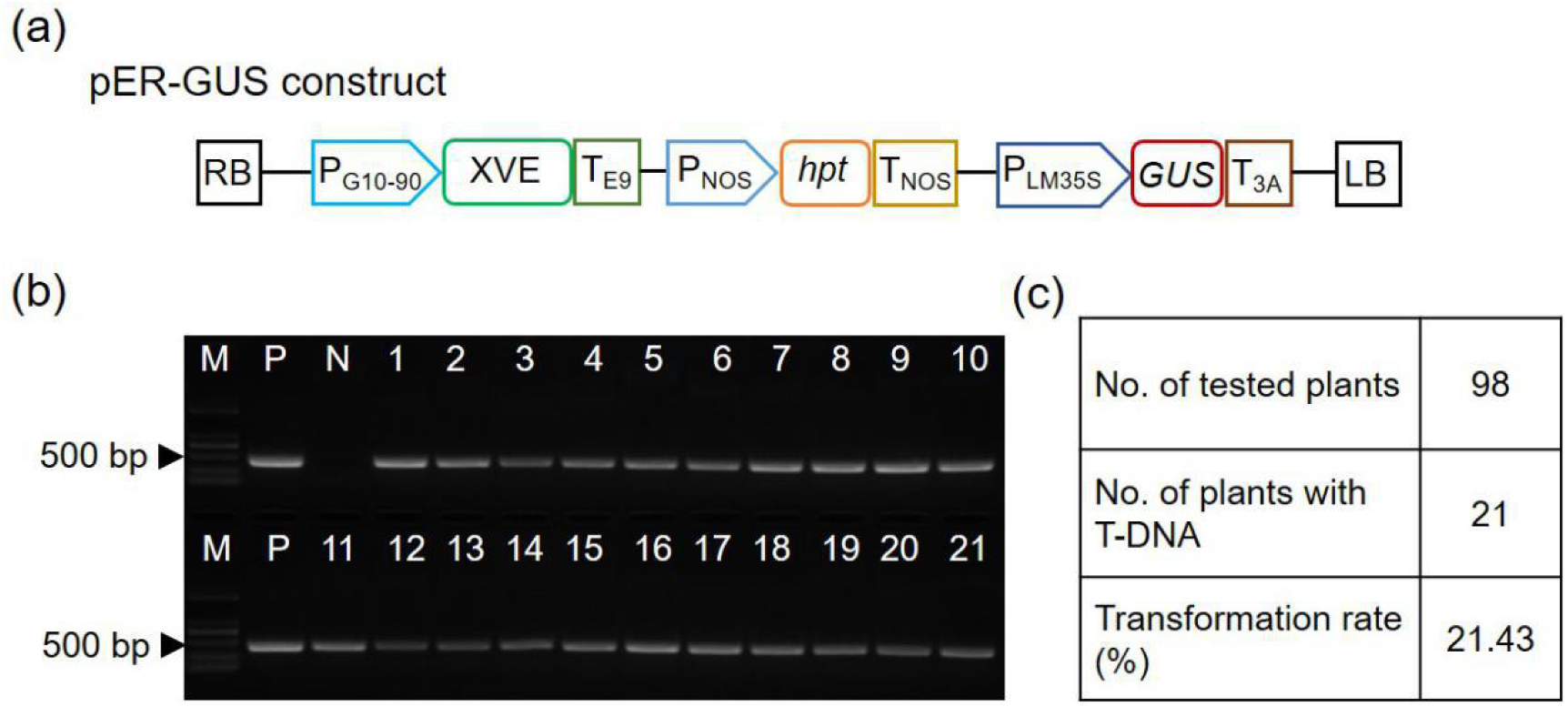
Generation of pER-GUS transgenic tobacco plants. (**a**) Schematic illustration of pER-GUS construct. P_G10-90_, G10-90 promoter; XVE, chimeric transactivator containing the regulator domain of an estrogen receptor; T_E9_, rbcS E9 terminator; P_NOS_, nopaline synthase promoter; htp, hygromycin phosphotransferase gene; T_NOS_, nopaline synthase terminator; P_LM35S_, 8 × LexA DNA binding site fused with the −46 CaMV 35S minimal promoter; GUS, β-glucuronidase; T_3A_, rbcS 3A terminator; RB, right border; LB, left border. (**b**) PCR identification of transgenic plants with *GUS*-specific primers. The plasmid and wild-type DNA were used as the positive control (P) and negative control (N), respectively. M, DNA marker; NC, no template control; lanes 1-21, different plant lines. (**c**) Overview of the identification of transgenic plants..

**Figure 2.**
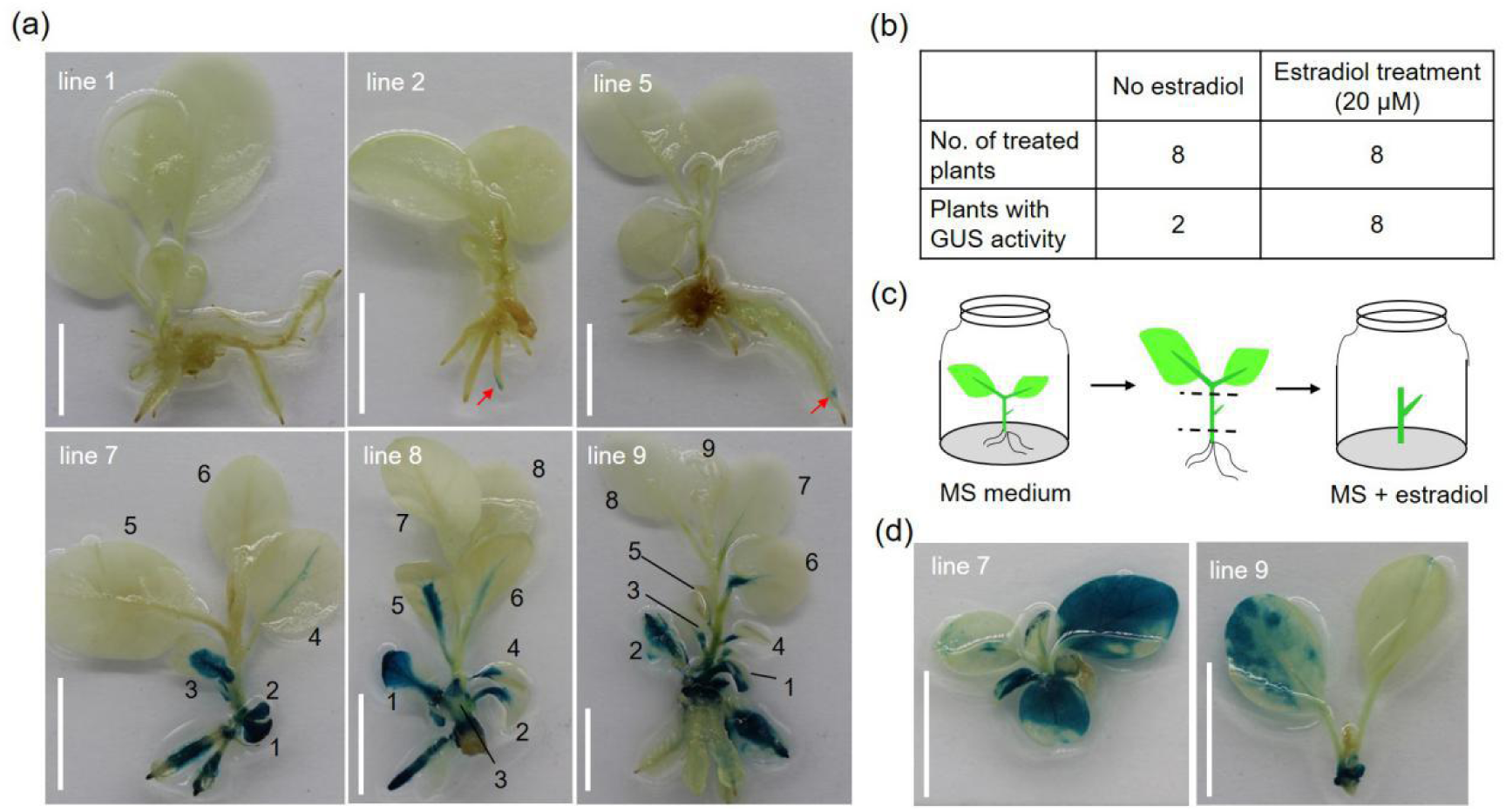
Histochemical staining assay of pER-GUS transgenic plants. (**a**) GUS staining of pER-GUS transgenic plants treated with 0 or 20 µM estradiol. The self-rooted plants after subculture were divided into two groups for estradiol treatment. The leaves were numbered consecutively from the base of the treated plants (lower panel). The weak GUS staining observed in plants treated with 0 µM estradiol (upper panel) was indicated in red arrows. Scale bars = 1 cm. (**b**) Overview of GUS staining results. The small stems with single axillary buds derived from subcultured plants (**c**) were used for estradiol treatment, and GUS staining results were shown in (**d**). Scale bars = 1 cm.

The results of histochemical staining assay suggest that systemic movement of estradiol was limited within tobacco plants (Figure 2a). We therefore used small stem with an axillary bud for induction analysis (Figure 2c). Stems derived from line 7 and 9 were cultured on induction medium for four weeks, and GUS activity was then determined by histochemical staining. For transgenic line 7, GUS staining was detected in all the leaves of the newly developed plant. Similar results was observed in transgenic line 9, despite the difference in GUS staining intensity between different leaves (Figure 2d). Taken together, the obtained results showed that XVE could regulate the expression of *GUS* reporter gene stringently and efficiently in transgenic tobacco plants upon estradiol treatment.

### 2.2. XVE-mediated targeted mutagenesis in transgenic tobacco plants

To develop the inducible CRISPR/Cas9 system, the *Cas9* gene was amplified from pCACRISPR/Cas9 vector [10] and was cloned into pER8 instead of *GUS* reporter gene (Figure 3a). Given that knockout of *PDS* gene generally leads to visible albino phenotype [11,37], a single guide RNA (sgRNA) targeting the *NtPDS* gene [39] was introduced into pER-Cas9 construct to generate the final expression vector pER-Cas9-NtPDS, in which the sgRNA is driven by AtU6 promoter (Figure 3a). After transformation, a number of 15 plants were identified as transgenic lines after hygromycin-dependent screening, followed by PCR identification with *hpt*-specific primers (Figure 3b, c). The amplified *hpt* gene fragments were further verified by Sanger sequencing (Figure S4). Surprisingly, three lines (line #5, #7 and #13) showed incomplete pale phenotypes without estradiol treatment (Figure 3d), and targeted mutagenesis was observed at the target sites (data not shown), suggesting the “leaky” expression of *Cas9* in these plants. In addition, the development of transgenic plants carrying pER-Cas9-NtPDS expression cassette appeared less affected when compared with wild-type plants (Figure 3d).

**Figure 3.**
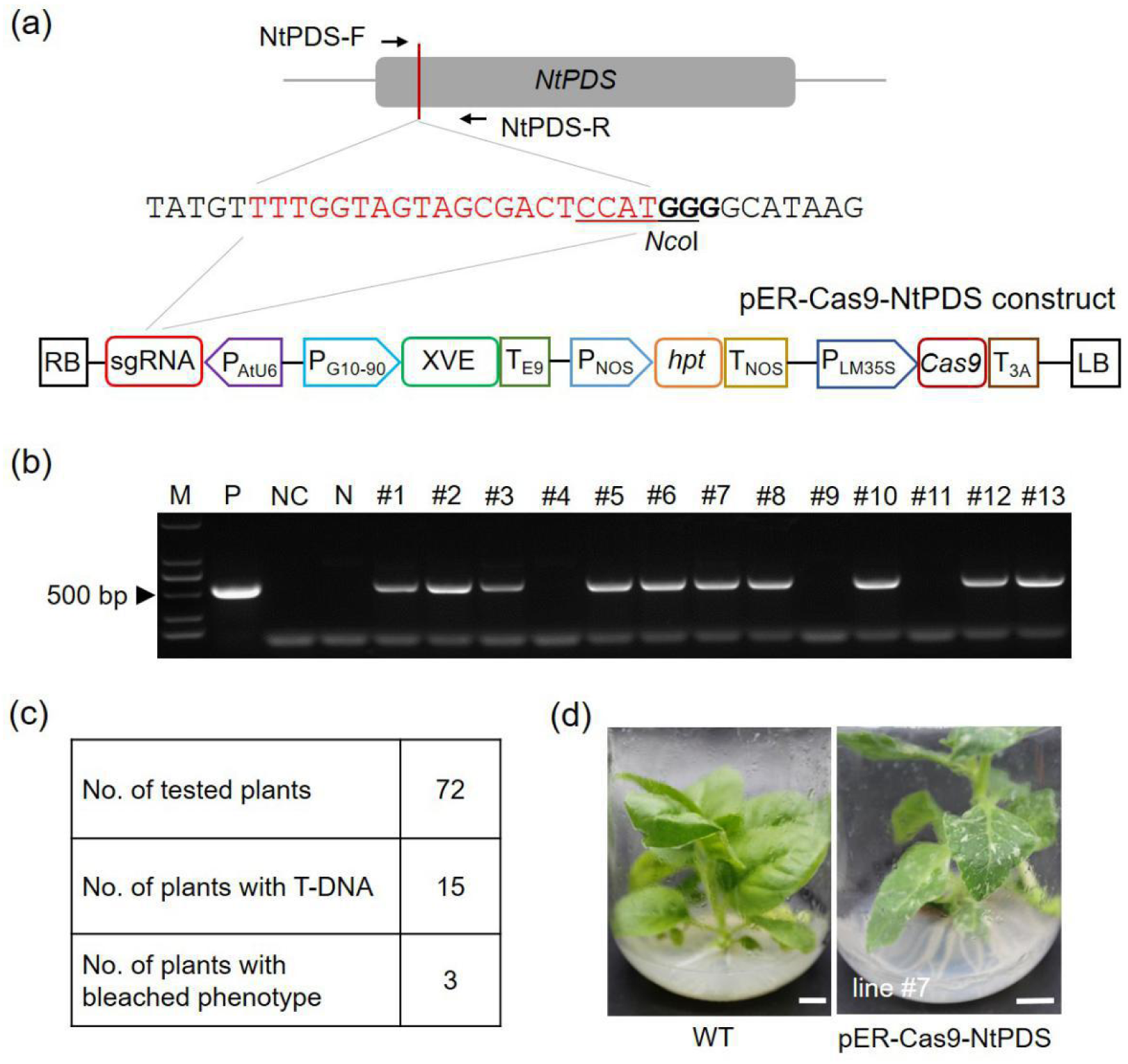
Generation of pER-Cas9-NtPDS transgenic tobacco plants. (**a**) The target sequence of *NtPDS* gene and schematic diagram of pER-Cas9-NtPDS construct. The 20-bp target sequence is indicated in red, and the PAM (protospacer adjacent motif) is in bold. The recognition sequence of *Nco*I restriction enzyme is underlined. sgRNA, single guide RNA; Cas9, CRISPR-associated protein 9. (**b**) PCR identification of transgenic plants using *hpt*-specific primers. The plasmid and wild-type DNA were used as the positive control (P) and negative control (N), respectively. M, DNA marker; NC, no template control; lanes #1-#13, different plant lines. (**c**) Overview of PCR identification results. (**d**) Phenotype of transgenic plant with leaky expression of *Cas9*. The pER-Cas9-NtPDS transgenic line #7 with leaky expression of *Cas9*, as well as wild-type (WT) regenerated plant, is shown. Scale bars = 1 cm.

Four independent transgenic lines (line #1, #2, #6 and #10) without phenotypic alterations were selected for induction experiment. One-week-old plants after subculture were transferred to the estradiol-containing (20 µM) MS medium and were cultured for another four weeks. After estradiol treatment, the basal leaves of all the four transgenic plants turned pale green (Figure 4a), suggesting disruption of the *NtPDS* gene. To investigate whether the phenotypic changes are caused by induced genome editing, we first checked the target sites in the four tested transgenic lines before estradiol treatment. The restriction enzyme (RE)/PCR assay is a useful method for mutation identification, and mutated genomic DNA, which is recalcitrant to enzyme digestions due to the destruction of available restriction enzyme sites, can be amplified by PCR [40,41]. The RE/PCR assay was performed with genomic DNA prepared from the four lines. The PCR results showed that no desired band was produced using *Nco*I-digested genomic DNA, similar to that of wild-type genomic DNA (Figure 4b). Additionally, the target region of *NtPDS* was amplified from the four lines, respectively, and was analyzed by Sanger sequencing, considering that Sanger sequencing assay of PCR products can directly uncover mutations from a pool of DNA sequences [42]. The sequencing chromatograms of the four lines turned out to be clear peaks (Figure 4c), suggesting that the target sequences in the four transgenic lines are unedited. Altogether, our sequencing results (Figure S5), together with RE/PCR results, demonstrated that there was no mutation at the target site in *NtPDS* gene in the four transgenic lines before estradiol treatment.

**Figure 4.**
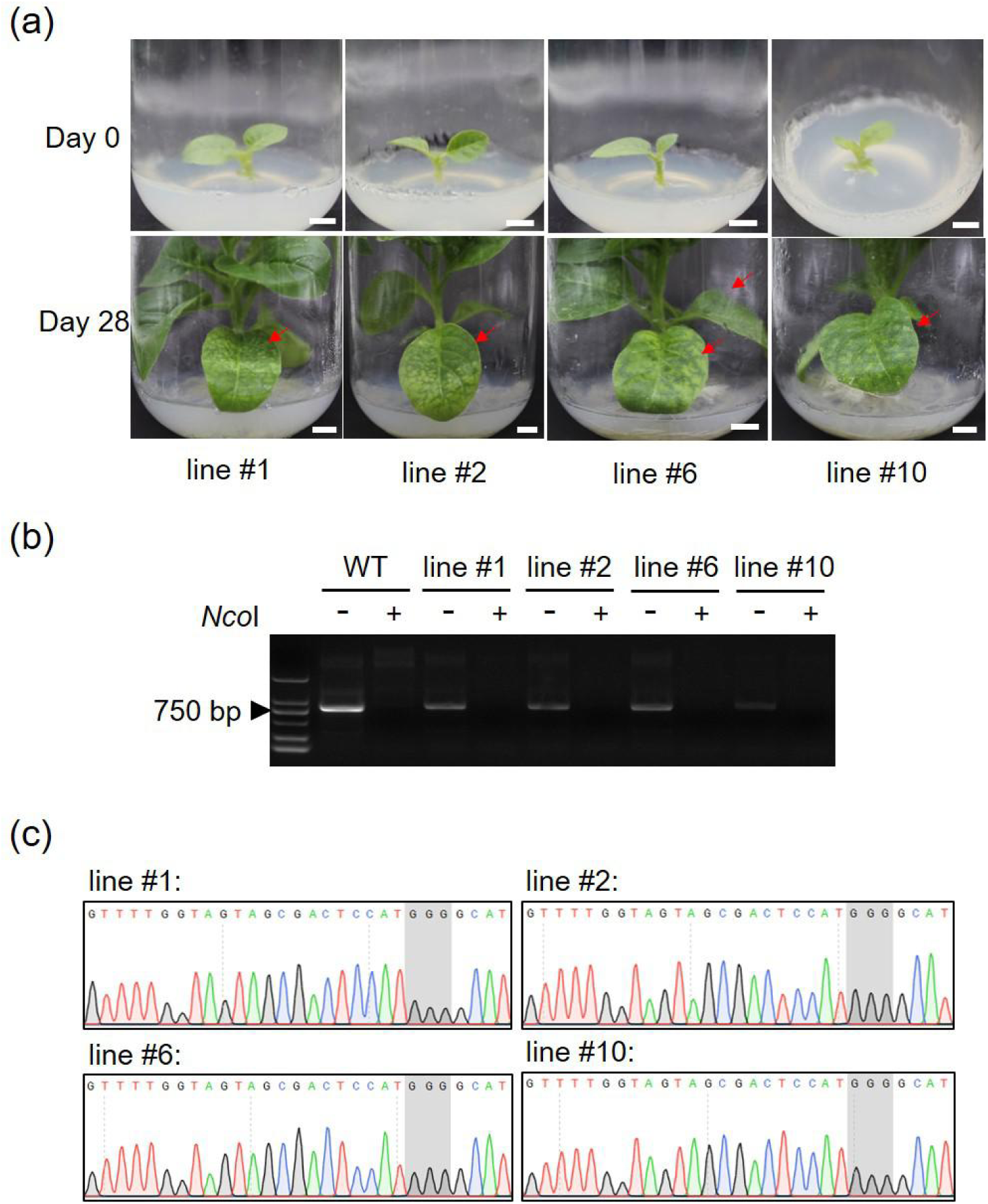
Verification of the target site in *NtPDS* gene in pER-Cas9-NtPDS transgenic plants before estradiol treatment. (**a**) The transgenic plantlets before (Day 0) and after (Day 28) estradiol treatment. Four independent transgenic lines, line #1, #2, #6 and #10, were used for treatment. One-week-old plants after subculture (Day 0) were transferred to estradiol-containing (20 µM) medium and were cultured for another 4 weeks (Day 28). The excised leaves of the four plantlets before estradiol treatment were sampled for subsequent identification. The pale green leaves developed after estradiol treatment were indicated in red arrows. Scale bars = 1 cm. (**b**) Restriction enzyme (RE)/PCR assay. The genomic DNA prepared from untreated plants was digested with *Nco*I enzyme to remove wild-type (WT) DNA. Only mutated sequences with destructed restriction enzyme site are recalcitrant to digestion and can be amplified by PCR. +, digestion with *Nco*I; -, no digestion. (**c**) Chromatograms of Sanger sequencing. The target sequences amplified with digested or undigested genomic DNA were analyzed by Sanger sequencing. The wild-type or unedited sequences generated sequencing chromatograms with single peaks [42]. The PAM sequences are highlighted in grey.

We then sampled the pale green leaves (Figure 5a) of the four transgenic lines after estradiol treatment for subsequent analysis. Similarly, we conducted RE/PCR assay, and desired bands, however, were produced using *Nco*I-treated genomic DNA (Figure 5b), suggesting the presence of mutations in genomic DNA sequences. Moreover, Sanger sequencing results of PCR products revealed targeted mutagenesis in *NtPDS* gene (Figure 5c,d). Targeted mutagenesis resulted in overlapping peaks starting from the mutation site (Figure 5c), which is in agreement with previous report [42]. Sequencing results of PCR amplicons uncovered indel mutations at the target site. Most mutations were single nucleotide insertions or deletions (Figure 5d), which is consistent with previous reports in other plant species [9,43]. Notably, at least two types of mutations, as well as wild-type sequences, were detected in each line (Figure 5d), suggesting that these transgenic plants were probably chimeras. Taken together, these results showed that *NtPDS* gene in the four transgenic lines was successfully edited after estradiol treatment, and mutaion of *NtPDS* resulted in development of pale green leaves.

**Figure 5.**
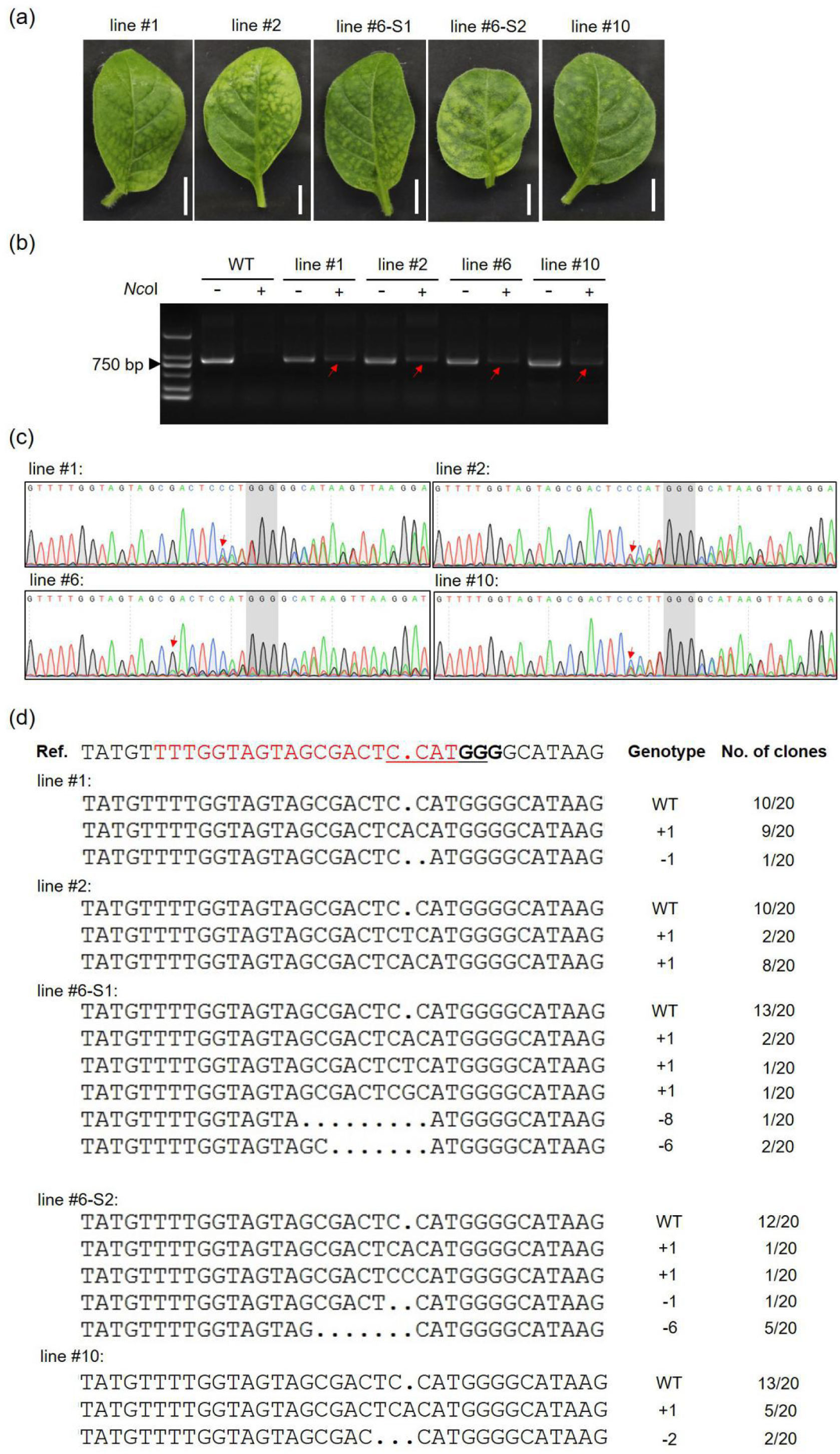
Induced mutation of *NtPDS* gene in pER-Cas9-NtPDS transgenic plants after estradiol treatment. (**a**) The leaves used for mutation detection. The pale green leaves of transgenic line #1, #2, #6 and #10 were sampled for our analysis. The two leaves of transgenic line #6 were marked as line #6-S1 and line #6-S2, respectively. Scale bars = 1 cm. (**b**) RE/PCR assay. The genomic DNA was digested with *Nco*I enzyme and was then used for PCR amplification. The bands of possible mutated DNA sequences are indicated in red arrows. +, digestion with *Nco*I; -, no digestion. (**c**) Sequencing chromatograms of the target sequences. The PCR products were directly analyzed by Sanger sequencing. Overlapping peaks were produced starting from the mutation sites (indicated in red arrows) near the PAM sequences (highlighted in grey) in the sequencing chromatograms [42]. (**d**) Mutation types identified from each transgenic line. The PCR products were further cloned into pLB vector and 20 clones of PCR amplicons were analyzed by Sanger sequencing. The mutation types and corresponding number of clones are shown on the right. The sequence of sgRNA is in red, and the PAM sequence is in bold. The recognition sequence of *Nco*I restriction enzyme is underlined. Ref., reference sequence.

### 2.3. The expression of Cas9 is strongly induced by estradiol

The expression profiles of *Cas9* in the four tested transgenic lines (line #1, #2, #6 and #10) with or without estradiol treatment were characterized using qRT-PCR assay. As mentioned above, three independent lines (line #5, #7 and #13) exhibited incomplete albino phenotypes without estradiol treatment (Figure 3d), and qRT-PCR results revealed a relatively high expression level of *Cas9* in these plants in the absence of exogenous estradiol (Figure 6). By contrast, the transcript level of *Cas9* in line #1, #2, #6 and #10 without estradiol treatment was extremely low when compared with the leaky expression lines (Figure 6). However, after exposure to estradiol, the transcript abundance of *Cas9* in the four transgenic lines was significantly increased (Figure 6), suggesting that the expression of *Cas9* is strongly induced by exogenous estradiol.

**Figure 6.**
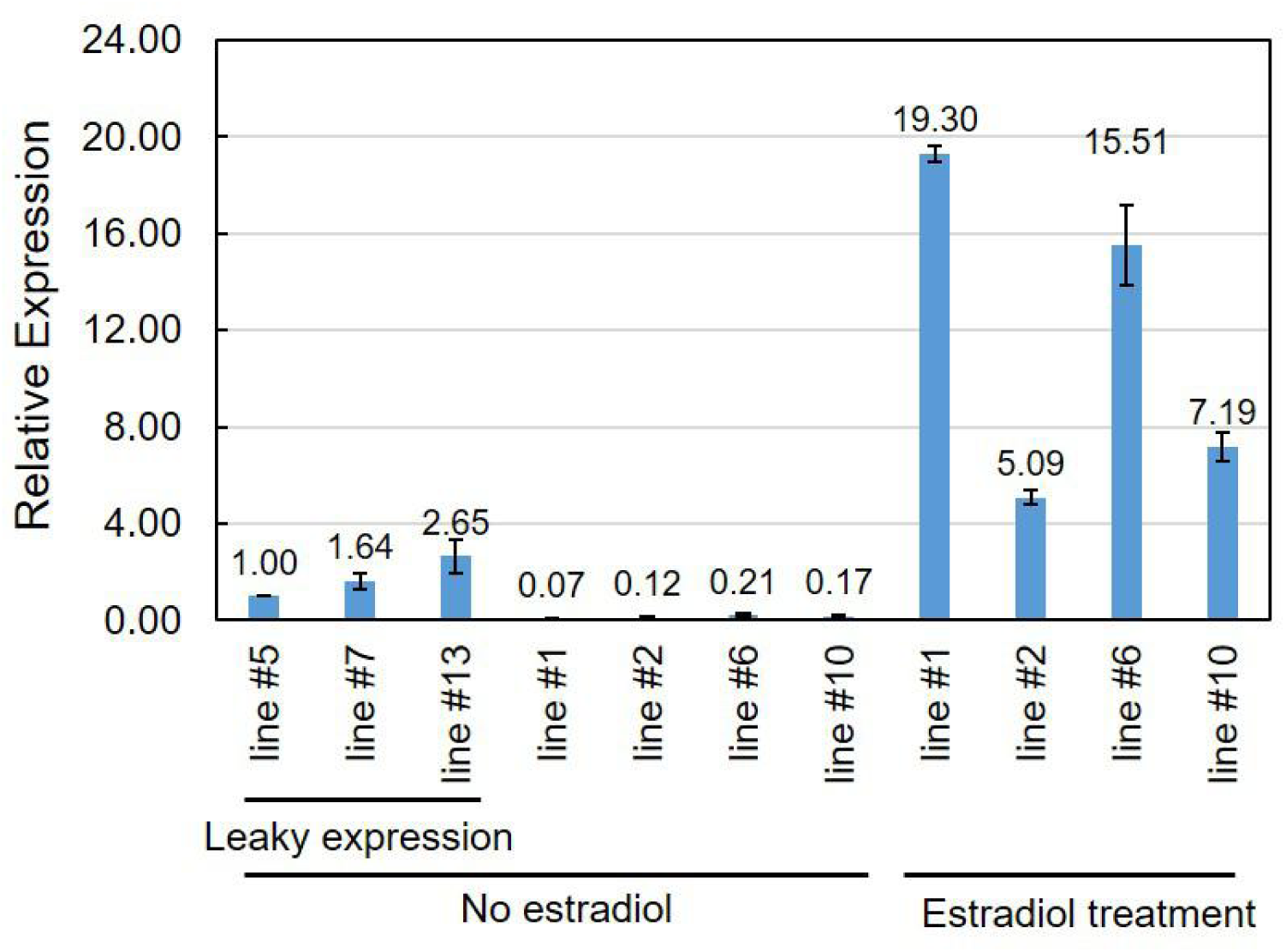
Expression profiles of *Cas9* gene in pER-Cas9-NtPDS transgenic lines upon estradiol treatment. The transgenic line #5, #7 and #13 with white parts in their leaves under uninduced conditions had leaky expression of *Cas9*. The line #5 was used as the control and the expression of *Cas9* in other transgenic lines relative to line #5 was determined by qRT-PCR. The relative expression level of *Cas9* was calculated using the 2 ^− Δ Δ CT^ method [51] with the tobacco β-Tubulin encoding gene (accession number: U91564) being used as the internal control. The experiment was repeated three times. The data is shown as means ± SD.

### 2.4. The type of explants used for estradiol treatment has an effect on genome editing

According to the GUS staining results described above, the use of single axillary bud for treatment with estradiol appeared to contribute to strong GUS staining in developed plants (Figure 2d). Thus, we used small stems with single axillary buds of transgenic line #1 and #2 for induction analysis. After treatment with estradiol, the newly developed plantlets showed white rather than pale green parts in their leaves (Figure 7a), suggesting knockout of *NtPDS* gene in these plants. As expected, indel mutations were detected in *NtPDS* gene in the two transgenic lines (Figure 7b).

**Figure 7.**
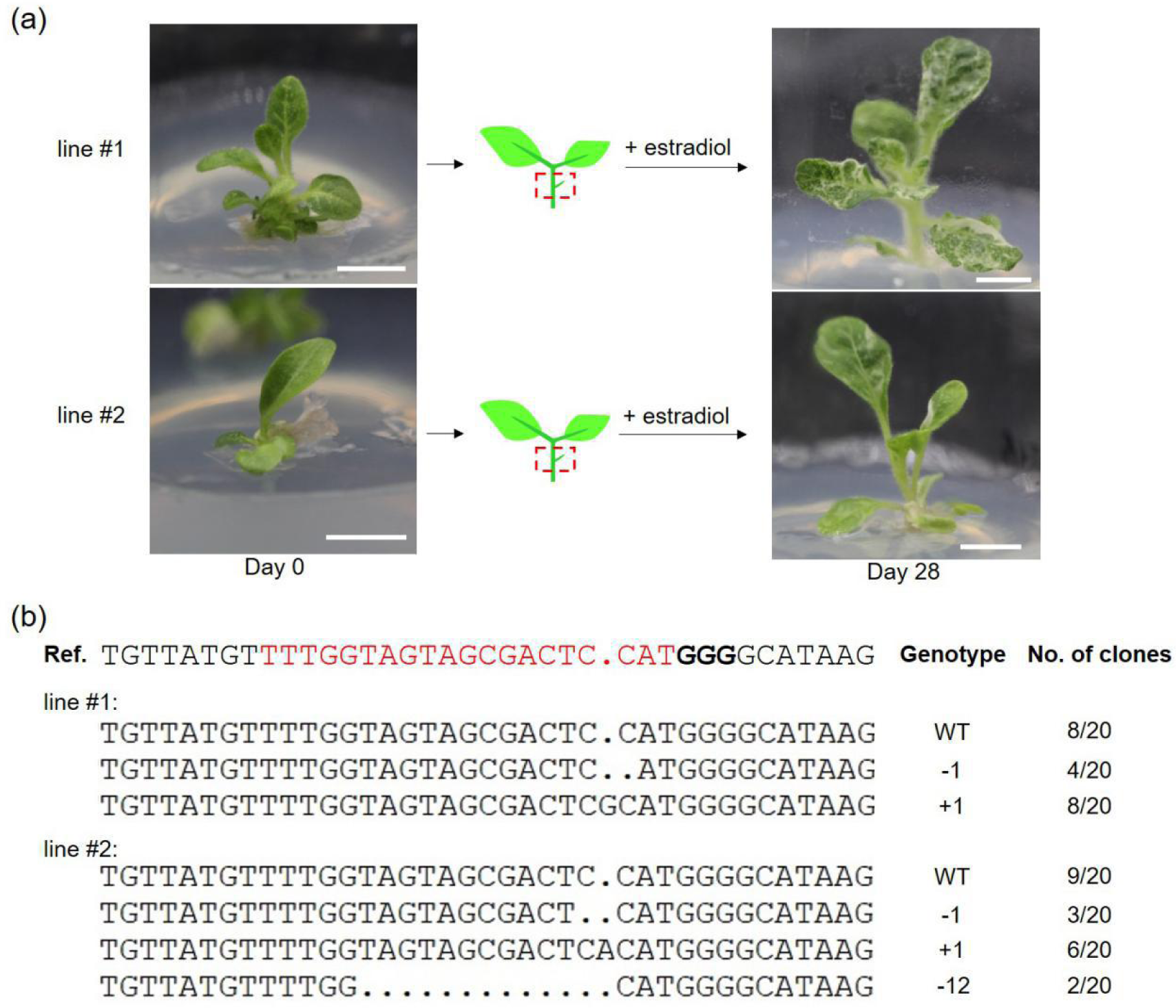
Induction analysis with single axillary buds. (**a**) Estradiol treatment of single axillary buds. The leaves and roots of in vitro plantlets (Day 0) were excised and small stems with single axillary buds (indicated in red rectangles) were retained and used as explants for estradiol treatment. Scale bars = 1 cm. (**b**) Mutations detected in estradiol-treated plants shown in (**a**). The mutation types and corresponding number of clones are shown on the right. The sequence of sgRNA is in red, and the PAM sequence is in bold.

## 3. Discussion

In general, plant genome editing is conducted using the CRISPR/Cas9 system, in which the *Cas9* gene is usually driven by a constitutive CaMV 35S promoter or plant native promoter [10,44,45,46]. However, constitutive expression of *Cas9* is not necessary, given the fact that transient expression of Cas9 in plants is enough to induce the targeted mutagenesis [35,45,46]. Moreover, knockout of fundamentally significant genes (almost 10% of the Arabidopsis genome) would result in severe pleiotropic effects or lethality [47,48], and the lack of corresponding loss-of-function mutants makes it impossible to decipher the functions of these indispensable genes. More accurate editing would be preferred considering that many genes express in specific context in plants [48]. Inducible genome editing can be used as an approach to overcome the limitations and accomplish temporal and spatial editing of genes of interest. The development of inducible genome editing system would expand the CRISPR toolbox and allows us to perform genome editing more flexibly. In the present study, we developed an estradiol-inducible CRISPR/Cas9 system and conducted genome editing in *N. tabacum* using this inducible system by targeting *NtPDS* gene (Figure 5 and 7). Both little plantlets and small stems with single axillary buds were used for estradiol treatment (Figure 4a and Figure 7a). The plantlets developed pale green leaves (Figure 4a) while the newly generated plants developed from buds showed white parts in leaves after estradiol treatment (Figure 7a), suggesting that the type of explants could affect the genome editing. In fact, in previous report, expression pattern of *GUS* reporter gene upon estradiol treatment varied in different tissues or organs [19]. In addition, all the tested lines (line #1, #2, #6 and #10) were found to be chimeric after induced genome editing, and several mutation types, as well as wild-type sequences, were identified from each transgenic line (Figure 5 and 7). The primary reason is that we used the T0 plants for estradiol treatment, and it is nearly impossible for the edited cells to develop into new plants. Alternatively, transgenic seeds could be better materials for induced genome editing to generate edited plants. However, generation of chimeric plants avoids the adverse effect of knockout of those developmentally important genes, and transgenic plants with genetic chimera can serve as important materials for study of signaling mechanism in plants [31].

The results of qRT-PCR showed that the expression level of *Cas9* upon estradiol treatment varied in different transgenic lines (Figure 6), and the transcript abundance of *Cas9* can also affect the editing efficiency [49]. Furthermore, the expression level of XVE in transgenic plants also contributes to induction level of gene of interest [32]. In the XVE construct pER8, a weak constitutive G10-90 promoter was used to drive the expression of XVE [18,32,50]. Moreover, the difference of estradiol uptake between different plants had an effect on induction of target gene [32]. In fact, histochemical staining assay with transgenic lines harboring pER-GUS expression cassette revealed the difference in staining intensity bewteen different lines (Figure 2a and Figure S3). The other factors, such as the inserting loci and copies of T-DNA, may affect the induction level of target gene as well. Intriguingly, clear GUS staining was mainly detected in roots and lower leaves (Figure 2a). The limited movement of estradiol within the plants could be the reason why the leaves localized at the base of plants turned pale green after estradiol treatment (Figure 4a). The concentration of estradiol and induction time are another two factors that should be considered during induction. Though the XVE system can be efficiently activated by a low concentration (8 nM) of estradiol, varying inducer concentrations were reported in different plant species [16,18,19,35]. In Arabidopsis, a relatively low concentration (∼5 µM) of estradiol was used for treatment [18,19], and in rice, however, the inducer concentration increased to 20 µM [16,35]. A long incubation resulted in reduced transcript level of target gene due to instability of the inducer [18]. However, expected phenotypes were still observed in transgenic Arabidopsis plants after a prolonged incubation (29 d) in the presence of the inducer [19]. Moreover, transient expression of *Cas9* is sufficient to generate targeted mutagenesis [35,45]. Considering the low inducibility of target gene in leaves of intact transgenic plants treated with estradiol through root absorption [16,32], the explants used in this study were therefore cultured on estradiol-containing (20 µM) medium for a long time (28 d). Optimization of the inducer concentrations and induction time, of course, would help to improve the expression of *Cas9* upon induction.

Three transgenic lines (line #5, #7 and #13) with white parts in their leaves (Figure 3d), as well as the four test lines (line #1, #2, #6 and #10), were observed to have leaky expression of *Cas9* under uninduced conditions (Figure 6). The leaky expression of the XVE system observed here was also reported in previous studies [16,32,35]. However, the leaky expression effect is generally very weak, and our GUS staining assay with pER-GUS transgenic plants revealed faint expression of GUS in roots (Figure 2a). Actually, the expression level of *Cas9* in the line #1, #2, #6 and #10 was extremely low (Figure 6), and no mutation was detected in *NtPDS* gene in these plants without treatment with estradiol (Figure 4 and Figure S5).

In conclusion, the XVE-based CRISPR/Cas9 system appeared to be an effective tool for induced genome editing in tobacco, and it promises to be a useful approach that allows temporal and spatial control of genome manipulation in plants after further optimization.

## 4. Materials and Methods

### 4.1. Plasmid construction

The plant expression vector pER8 (GenBank: AF309825.2) was linearized by digestion with *Xho*I and *Spe*I (NEB). To develop the pER-GUS construct, the *GUS* gene was amplified from pBI121 (vector information is available in snapgene, https://www.snapgene.com/) using the KOD-Plus-Neo kit (TOYOBO) with GUS-pER-F and GUS-pER-R primers (Table S1). The PCR was conducted in a 50 µL volume and the procedure is as follows: 94 °C for 3 min; 32 cycles of 98 °C for 10s, 58 °C for 30s, and 68 °C for 130s, followed by a final extension at 68 °C for 5 min. The amplified *GUS* gene was cloned into linearized pER8 using the ClonExpress II one step cloning kit (Vazyme) via homologous recombination (HR). Similarly, the *Cas9* gene amplified from pCACRISPR/Cas9 [10] using Cas9-pER-F and Cas9-pER-R primers (Table S1) was cloned into pER8 to develop the pER-Cas9 vector. The AtU6-sgRNA expression cassette, which contains the sgRNA targeting *NtPDS* gene, was amplified using AtU6-sgRNA-F and AtU6-sgRNA-R primers (Table S1) from well-constructed Cas9-NtPDSg2 vector described in our previous study [39]. The AtU6-sgRNA expression cassette was inserted into the pER-Cas9 vector via the *Pme*I site by HR. The constructed vector was designated as pER-Cas9-NtPDS.

**Table S1.**
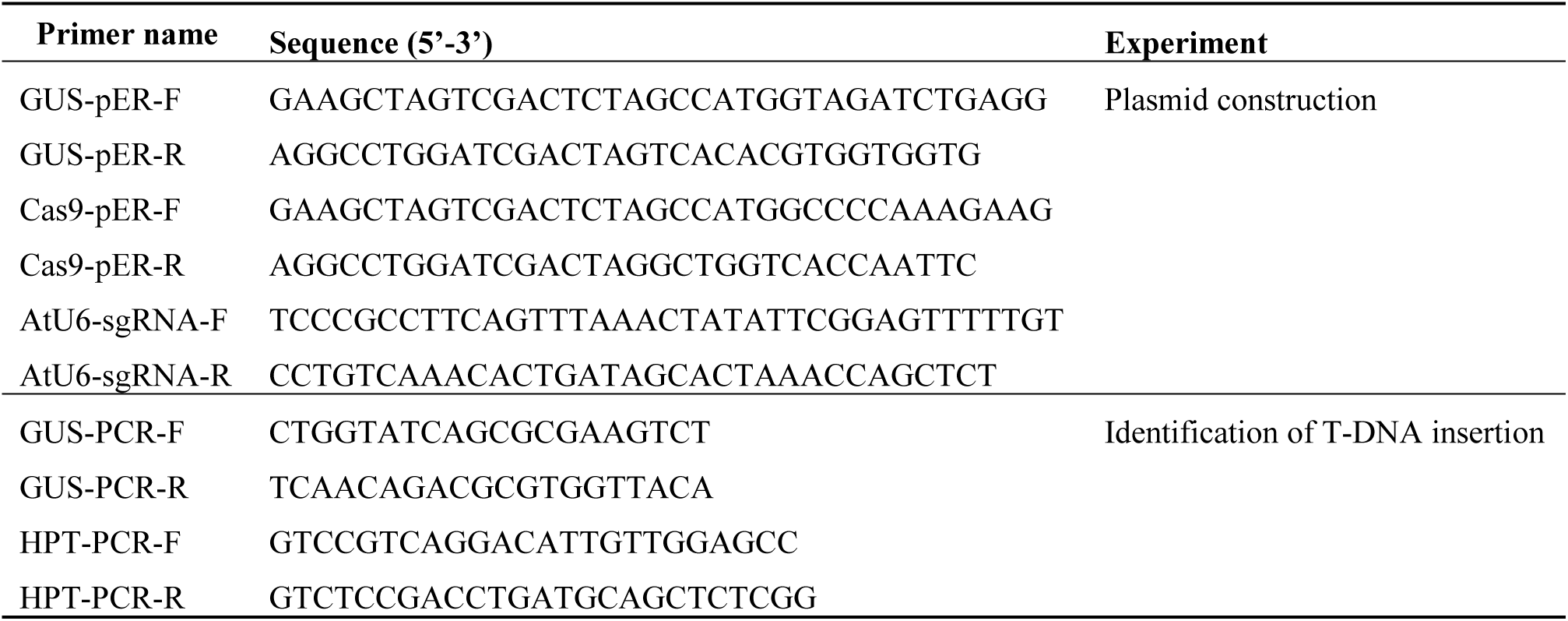
List of primers used in this study.

### 4.2. Plant transformation

The surface sterilized seeds of *N. tabacum* cv. Samsun germinated on Murashige and Skoog (MS) medium (PhytoTech), and one-month-old in vitro plants were used for transformation. The dissected tobacco leaves were incubated with *Agrobacterium tumefaciens* EHA105 cells, which contained the constructed plasmids. The transformation was performed as previously described [39].

### 4.3. Identification of transgenic plants

The leaves of regenerated plants were collected and genomic DNA was prepared using plant genomic DNA kit (Aidlab) following the manufacturer’s instructions. The specific primers designed for *GUS* or *hpt* gene were used for identification of transgenic plants. The PCR with a volume of 50 µL was carried out using Taq polymerase (Vazyme) at 95 °C for 3 min; 30 cycles of 95 °C for 15s, 60 °C for 15s, and 72 °C for 10s, followed by a final extension at 72 °C for 5 min. Around 10 µL of the PCR products were used for electrophoresis on an ethidium bromide-stained agarose gel (1%), and the remaining products were further verified by Sanger sequencing. The identified T0 plants were subcultured on MS medium and were used for estradiol treatment.

### 4.4. Estradiol treatment

The estradiol treatment was performed as previously described [16] with some modifications. A stock of 20 mM 17-β-estradiol (Sigma-Aldrich) in dimethyl sulfoxide (DMSO) was prepared and stored at −20 °C. The stock was added in solid MS medium at a final concentration of 20 µM before use. For estradiol treatment, one-week-old plants were transferred to MS medium with 20 µM estradiol and were cultured for four weeks. To conduct treatment of short stems with single axillary buds, the leaves and roots of in vitro plants were excised, and small stems with only one axillary bud were retained and used for estradiol treatment. The leaves of tobacco plants before and after estradiol treatment were sampled for genomic DNA and total RNA extraction, respectively.

### 4.5. Mutation detection

For RE/PCR assay, 500 ng of the genomic DNA was first treated with *Nco*I High-Fidelity restriction enzyme (NEB) overnight at 37 °C according to the manufacturer’s protocol, and then the reaction was incubated at 80 °C for another 20 min for heat inactivation. PCR was performed with KOD polymerase using 1 µL of the mixture as the template. The untreated genomic DNA was used as the control.

For Sanger sequencing assay, the prepared genomic DNA was directly used as the template for PCR. The PCR products were directly sent for Sanger sequencing or were further cloned into the pLB vector (TIANGEN). A number of 20 amplicons clones from each plants were analyzed by Sanger sequencing. All the PCR reactions were carried out with NtPDS-F and NtPDS-R primers as our previously described [39].

### 4.6. qRT-PCR analysis

The qRT-PCR assay was performed as described in our previous report [39]. Briefly, total RNA was prepared with HiPure HP plant RNA mini kit following the manufacturer’s instructions, and the first strand of cDNA was synthesized using HiScript III 1^st^ strand cDNA synthesis kit (Vazyme). The relative expression level of *Cas9* gene was measured by qRT-PCR using the 2^−ΔΔCT^ method [51] with the β-Tubulin encoding gene (accession number: U91564) being used as the internal control.

### 4.7. Histochemical staining

Histochemical staining was conducted using X-Gluc as substrate. The transgenic plants treated with 0 or 20 µM estradiol were immersed in GUS staining solution (Coolabler) and put in a vacuum equipment. The vacuum was kept at 0.085 MPa for 30 min. Then the samples were incubated overnight at 37 °C. After staining, the samples were cleared by 95% ethanol.

## Supplementary Materials

Supplementary materials are included in this study.

## Author Contributions

C.R., S.L. and Z.L. conceived the study; C.R., Y.L., X.W. and Y.G. performed the experiments; C.R. and Z.L. wrote the manuscript; P.F. and S.L. revised the manuscript.

## Acknowledgments

We thank Elias Kirabi Gathunga (Institute of Botany, the Chinese Academy of Sciences) for proof reading.

