## Supplementary material for "Targeted genome editing in *Nicotiana tabacum* using inducible CRISPR/Cas9 system": Supplemenal materials

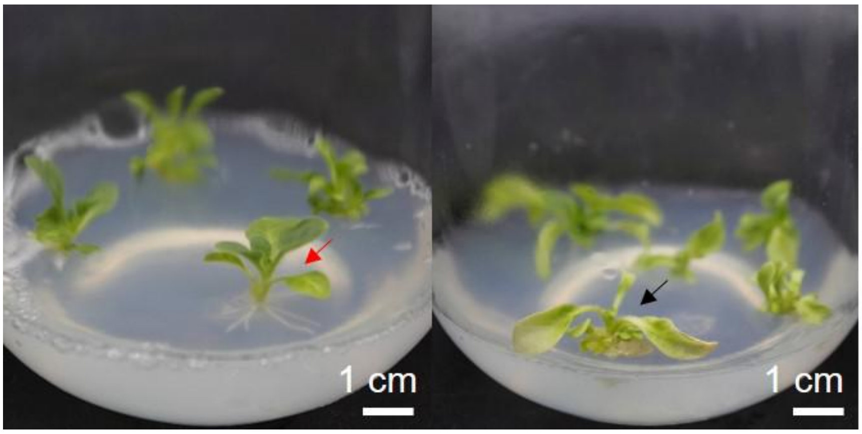


Figure S1. Screening of hygromycin-resistant plants. The plants that developed roots (indicated in red arrow) on hygromycin-containing medium were selected as candidates for PCR identification, while those plants that cannot develop roots (indicated in black arrow) were omitted from our analysis.

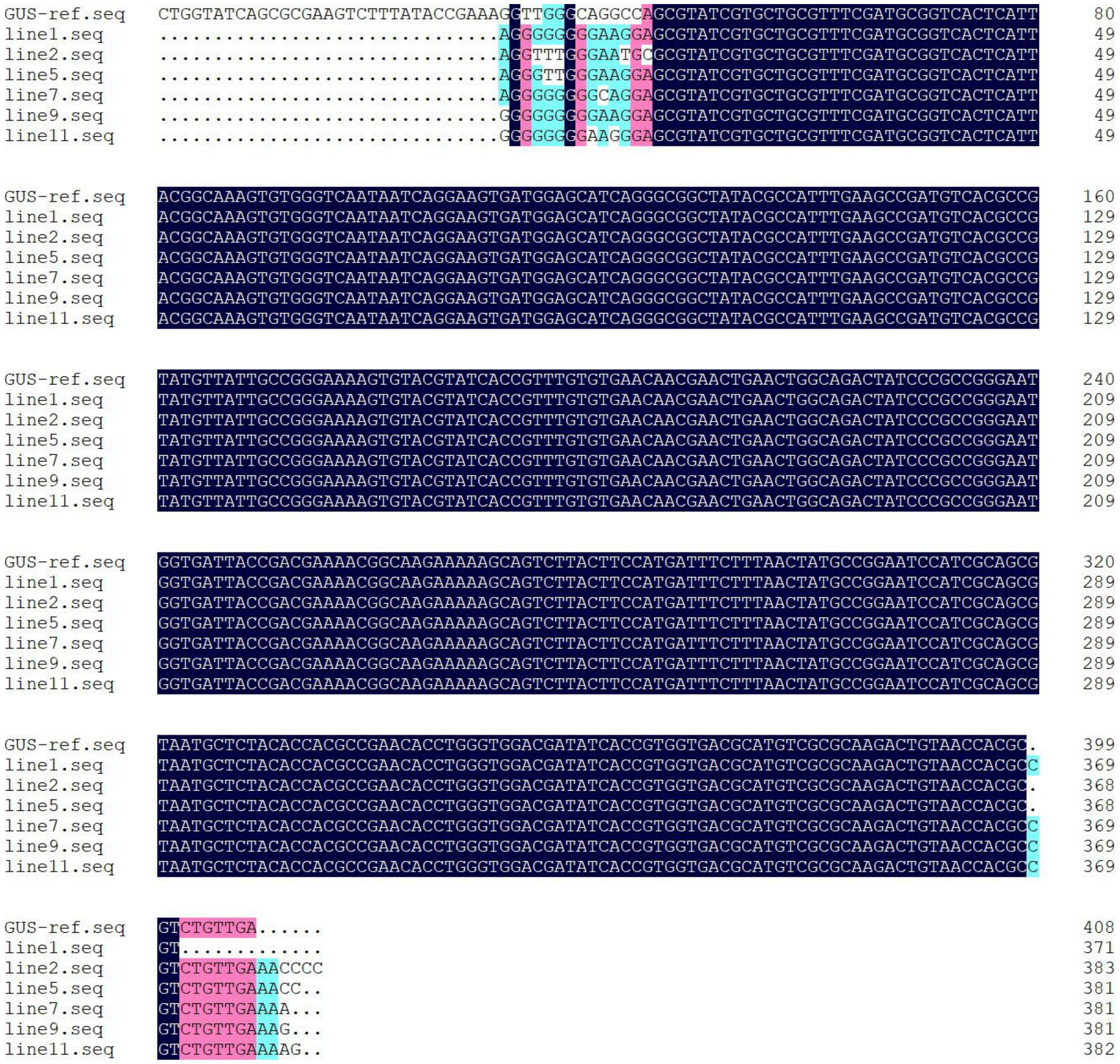


Figure S2. Sequencing result of the *GUS* reporter gene in pER-GUS transgenic plants. The gene fragments of *GUS* reporter gene were amplified from pER-GUS transgenic plants by PCR, and the products were verified by Sanger sequencing. The sequencing results of *GUS* gene fragments amplified from transgenic line 1, 2, 5, 7, 9 and 11 are shown.

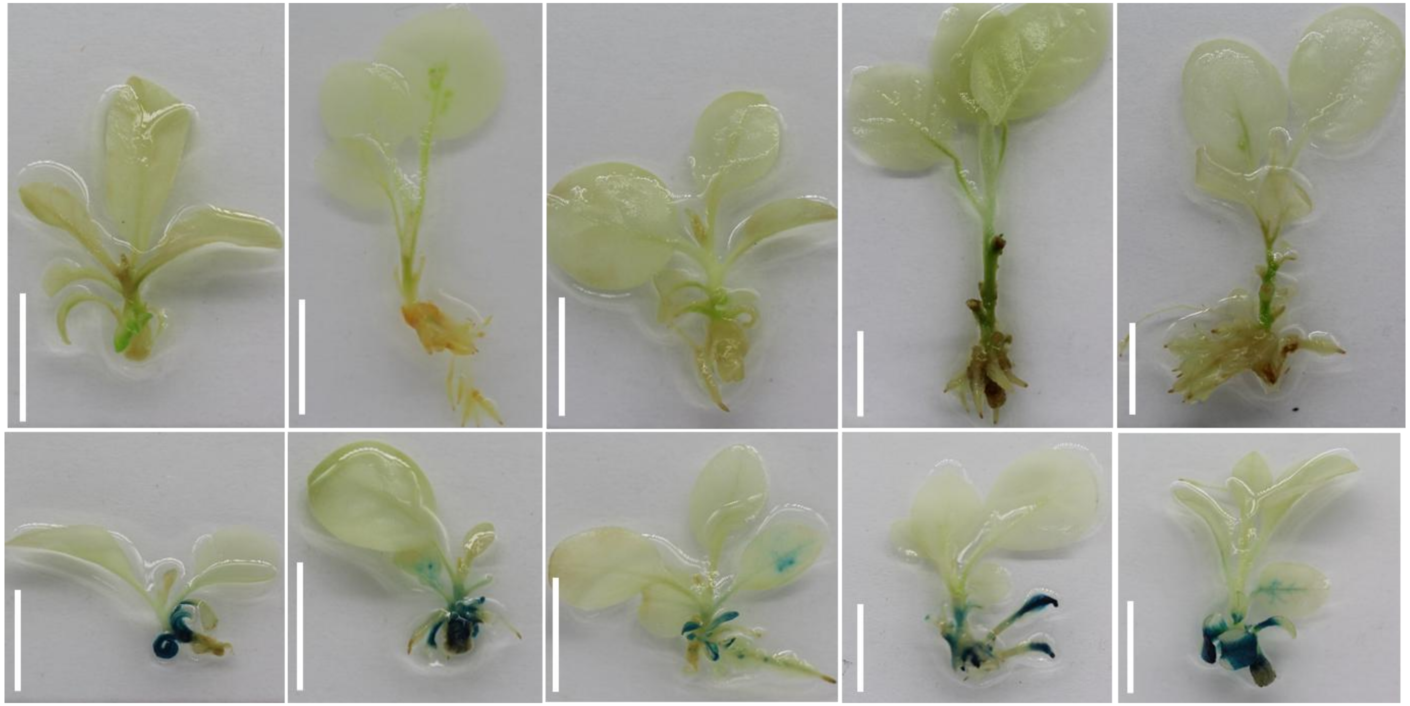

Figure S3. GUS staining of pER-GUS transgenic plants. The pER-GUS transgenic plants treated with 0 μM (upper pane) or 20 μM (lower pane) were sampled for GUS staining assay. Scale bars: 1 cm

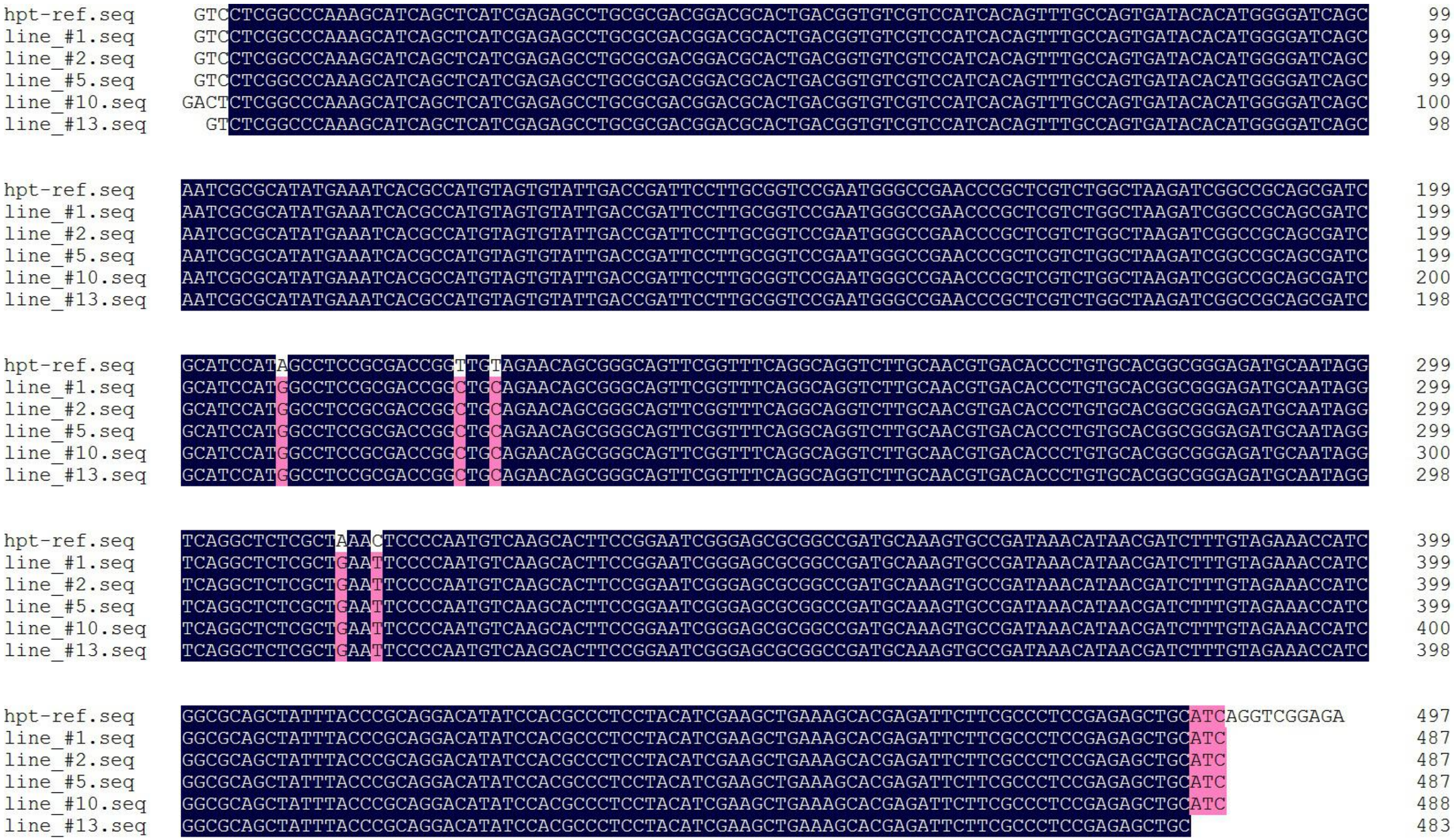

Figure S4. Sequencing result of the *hpt* resistant gene in pER-Cas9-NtPDS transgenic plants. The *hpt* gene fragments were amplified from transgenic plants and then verified by Sanger sequencing. The sequencing results of *hpt* gene fragments amplified from transgenic line #1, #2, #5, #10 and #13 are shown.

```
line_#1.seq        ATTTAGGTTCACAAGTGGGACAATCTTCTTACACTGAAATCAGGCTTAATTTACTGCTATTTTGTTCAGTAAAATGCCCCAAATTGGACTTGTTTCTGCC   199
line_#2.seq        ATTTAGGTTCACAAGTGGGACAATCTTCTTACACTGAAATCAGGCTTAATTTACTGCTATTTTGTTCAGTAAAATGCCCCAAATTGGACTTGTTTCTGCC   200
line_#6.seq        ATTTAGGTTCACAAGTGGGACAATCTTCTTACACTGAAATCAGGCTTAATTTACTGCTATTTTGTTCAGTAAAATGCCCCAAATTGGACTTGTTTCTGCC   200
line_#10.seq       ATTTAGGTTCACAAGTGGGACAATCTTCTTACACTGAAATCAGGCTTAATTTACTGCTATTTTGTTCAGTAAAATGCCCCAAATTGGACTTGTTTCTGCC   199
NtPDS-ref.seq                                                                             ATGCCCCAAATTGGACTTGTTTCTGCC    27
NtPDS-sgRNA.seq                                                                                                         0

line_#1.seq        GTTAATTTGAGAGTCCAAGGTAATTCAGCTTATCTTTGGAGCTCGAGGTCTTCTTTGGGAACTGAAAGTCAAGATGGTCACTTGCAAAGGAATTTGTTAT   299
line_#2.seq        GTTAATTTGAGAGTCCAAGGTAATTCAGCTTATCTTTGGAGCTCGAGGTCTTCTTTGGGAACTGAAAGTCAAGATGGTCACTTGCAAAGGAATTTGTTAT   300
line_#6.seq        GTTAATTTGAGAGTCCAAGGTAATTCAGCTTATCTTTGGAGCTCGAGGTCTTCTTTGGGAACTGAAAGTCAAGATGGTCACTTGCAAAGGAATTTGTTAT   300
line_#10.seq       GTTAATTTGAGAGTCCAAGGTAATTCAGCTTATCTTTGGAGCTCGAGGTCTTCTTTGGGAACTGAAAGTCAAGATGGTCACTTGCAAAGGAATTTGTTAT   299
NtPDS-ref.seq      GTTAATTTGAGAGTCCAAGGTAATTCAGCTTATCTTTGGAGCTCGAGGTCTTCTTTGGGAACTGAAAGTCAAGATGGTCACTTGCAAAGGAATTTGTTAT   127
NtPDS-sgRNA.seq                                                                                                         0

line_#1.seq        GTTTTGGTAGTAGCGACTCCATGGGGCATAAGTTAAGGATTCGTACTCCCAGTGCCATGACCAGAAGATTGACAAAGGACTTTAATCCTTTAAAGGTTTG   399
line_#2.seq        GTTTTGGTAGTAGCGACTCCATGGGGCATAAGTTAAGGATTCGTACTCCCAGTGCCATGACCAGAAGATTGACAAAGGACTTTAATCCTTTAAAGGTTTG   400
line_#6.seq        GTTTTGGTAGTAGCGACTCCATGGGGCATAAGTTAAGGATTCGTACTCCCAGTGCCATGACCAGAAGATTGACAAAGGACTTTAATCCTTTAAAGGTTTG   400
line_#10.seq       GTTTTGGTAGTAGCGACTCCATGGGGCATAAGTTAAGGATTCGTACTCCCAGTGCCATGACCAGAAGATTGACAAAGGACTTTAATCCTTTAAAGGTTTG   399
NtPDS-ref.seq      GTTTTGGTAGTAGCGACTCCATGGGGCATAAGTTAAGGATTCGTACTCCCAGTGCCATGACCAGAAGATTGACAAAGGACTTTAATCCTTTAAAGGTTTG   227
NtPDS-sgRNA.seq      TTTGGTAGTAGCGACTCCAT                                                                              20

line_#1.seq        TTTTGAATGCGGTGTGATTCTGAATTTATGATCTTGGGCATATATTCTCTAAAATAAGAGAAGTATATCTTGCCATTCAGGTAGTCTGCATTGATTATCC   499
line_#2.seq        TTTTGAATGCGGTGTGATTCTGAATTTATGATCTTGGGCATATATTCTCTAAAATAAGAGAAGTATATCTTGCCATTCAGGTAGTCTGCATTGATTATCC   500
line_#6.seq        TTTTGAATGCGGTGTGATTCTGAATTTATGATCTTGGGCATATATTCTCTAAAATAAGAGAAGTATATCTTGCCATTCAGGTAGTCTGCATTGATTATCC   500
line_#10.seq       TTTTGAATGCGGTGTGATTCTGAATTTATGATCTTGGGCATATATTCTCTAAAATAAGAGAAGTATATCTTGCCATTCAGGTAGTCTGCATTGATTATCC   499
NtPDS-ref.seq      TTTTGAATGCGGTGTGATTCTGAATTTATGATCTTGGGCATATATTCTCTAAAATAAGAGAAGTATATCTTGCCATTCAGGTAGTCTGCATTGATTATCC   327
NtPDS-sgRNA.seq

line_#1.seq        AAGACCAGAGCTAGACAATACAGTTAACTATTTGGAGGCGGCGTTATTATCATCATCATTTCGTACTTCCTCACGCCCAACTAAACCATTGGAGATTGTT   599
line_#2.seq        AAGACCAGAGCTAGACAATACAGTTAACTATTTGGAGGCGGCGTTATTATCATCATCATTTCGTACTTCCTCACGCCCAACTAAACCATTGGAGATTGTT   600
line_#6.seq        AAGACCAGAGCTAGACAATACAGTTAACTATTTGGAGGCGGCGTTATTATCATCATCATTTCGTACTTCCTCACGCCCAACTAAACCATTGGAGATTGTT   600
line_#10.seq       AAGACCAGAGCTAGACAATACAGTTAACTATTTGGAGGCGGCGTTATTATCATCATCATTTCGTACTTCCTCACGCCCAACTAAACCATTGGAGATTGTT   599
NtPDS-ref.seq      AAGACCAGAGCTAGACAATACAGTTAACTATTTGGAGGCGGCGTTATTATCATCATCATTTCGTACTTCCTCACGCCCAACTAAACCATTGGAGATTGTT   427
NtPDS-sgRNA.seq

line_#1.seq        ATTGCTGGTGCAGGTGATTTTTTCCAGTCATCTATATTTGTAGTCTTCATTTTTCTTTTTTCGGAGGGAAGATCATTCTATTAGTTGTATTATCACTAGA   699
line_#2.seq        ATTGCTGGTGCAGGTGATTTTTTCCAGTCATCTATATTTGTAGTCTTCATTTTTCTTTTTTCGGAGGGAAGATCATTCTATTAGTTGTATTATCACTAGA   700
line_#6.seq        ATTGCTGGTGCAGGTGATTTTTTCCAGTCATCTATATTTGTAGTCTTCATTTTTCTTTTTTCGGAGGGAAGATCATTCTATTAGTTGTATTATCACTAGA   700
line_#10.seq       ATTGCTGGTGCAGGTGATTTTTTCCAGTCATCTATATTTGTAGTCTTCATTTTTCTTTTTTCGGAGGGAAGATCATTCTATTAGTTGTATTATCACTAGA   699
NtPDS-ref.seq      ATTGCTGGTGCA                                                                                        439
NtPDS-sgRNA.seq
```

Figure S5. Analysis of the target sites in pER-Cas9-NtPDS transgenic line #1, #2, #6 and #10 before estradiol treatment. The target regions were amplified by PCR and verified by Sanger sequencing.
